# A first order approximation of a joint Potts and phylogenetic model

**DOI:** 10.64898/2026.09.22.753644

**Authors:** Remco Bouckaert

**Affiliations:** School of Computer Science, University of Auckland, Auckland, New Zealand

## Abstract

This study introduces a novel model that integrates the capacity of Potts models to capture structural dependencies with the computational efficiency afforded by Felsenstein’s algorithm in phylogenetics. While the present work focuses on protein models, the PhyloPotts framework is equally applicable to alignments of discrete states, including RNA, DNA, and 3DI characters. Our findings indicate that, although phylogenetic estimates do not consistently exhibit reduced bias relative to standard models, the approach substantially enhances the accuracy of ancestral sequence reconstruction.

The method is freely available via an open source implementation in the PhyloPotts package for BEAST 2 from https://github.com/rbouckaert/potts.

## 1 Introduction

Ignoring dependencies between sites in an alignment is a standard assumption in pretty much any popular phylogenetic software[1, 6, 8, 16, 27, 30]. The main attraction of this assumption is that it allows the likelihood of a tree for a given alignment to be calculated efficiently through Felsenstein’s peeling alogirhtm[7]. However, it is known that this assumption can result in biased tree estimates[12, 19].

Some attempts have been made to model dependencies between sites[12, 21, 24, 25, 34], going back some decades ago. There also exists specialist models, for example for single sequence systems designed for development biology[28]. None of these methods have been proven to be popular in a general setting.

Where phylogeneticists largely ignore dependencies between sites, structural biologists are *only* interested in those dependencies. The latter group starts with a sequence alignment and infer direct interactions between sites. In particular, Potts models[5, 10, 18] are popular for learning these interactions, and have been successfully applied in protein design[3], predicting 3D structures and protein contact[5], RNA structure prediction[4], to name just a few. Potts models are better at capturing structural interactions than autoregressive models[14] employed in [12]. Although methods exist weighing sequences based on their relatedness[11, 26], typically, the phylogeny representing the evolutionary history that created the sequences in the alignment is ignored.

Here, we combine Potts models with phylogenetic tree likelihood, thus allowing some dependencies between sites. This can be considered a combination of the directed graphical model representing the phylogentic likelihood with the undirected graphical model representing the Potts model. However, the Potts model only gouverns the root sequence, and internal node states still can be integrated out in a Felsenstein’s peeling algorithm like fashion. This means, we maintain the efficiency of standard phylogenetic methods, while at the same time capturing some of the dependencies between sites.

The methods is implemented in PhyloPotts package for BEAST2[1] in a Bayesian framework, and can be combined with any of the available tree priors, substitution models and clock models. This allows joint estimation of the tree topology and timing, any of the parameters involved in the tree prior, site model and clock model as well as ancestral reconstruction[9, 29] of sequences at the root and internal nodes of the tree.

## 2 Methods

First, we describe the Potts model and Felsenstein’s peeling algorithm separately. Then, we introduce a method to combine the two, and consider issues on how to simulate under the combined model.

### 2.1 Potts models

We assume we have an alignment of protein sequences, though note that the theory equally applies to any type of finite discretised sequence alignments, like RNA, DNA or 3DI[22]. A protein sequence of length *L* is treated as a system where each position *i* can exist in one of *q* = 21 states (20 amino acids states + 1 gap state). The model assigns a probability *P* (*S*) to any sequence *S* based on a *Hamiltonian (aka energy function)* defined as:

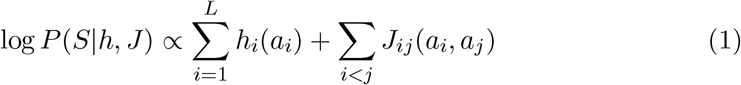

where *hi*(*ai*) represents the local preference for a specific amino acid at position *i* (similar to a amino acid profile, or position dependent stationary frequencies), and *Jij*(*ai, aj*) is the critical interaction term, which quantifies the direct statistical dependency between position *i* and position *j*.

There are an enormous number of parameters that need to be estimated, usually much more than the number of sites in the alignment. Joint estimation of Potts model and phylogeny requires dealing with the normalising constant (not shown in Eq(1)) that ensures that *P* (*S* | *h, J*) summed over all states is 1. This is computationally hard given there are 21*L* such states. Therefore, in this paper, we pre-estimate these parameters from a given alignment before inferring any phylogeny.

Though estimating the parameters *hi* and *Jij* can be computational prohibitive for large alignments, we only considered relatively small problems. Estimating the Potts model parameters through a Boltzman machine is the gold standard and we found it to be practical for smaller alignments. Note that due to the large number of parameters, there is a tendency for the model to overfit the data. To prevent this happening, regularisation was used in order to keep the magnitude of the parameters in check.

### 2.2 Felsenstein’s phylogenetic likelihood

Felsenstein’s phylogenetic likelihood is the standard statistical framework used to calculate the probability of observing a set of DNA or protein sequences given a specific evolutionary tree and a model of mutation. Felsenstein’s breakthrough was the *Pruning Algorithm*, a dynamic programming approach that turned an exponential problem into a linear one. To calculate the likelihood of a tree, one needsto know what happened at every internal node. However, we only observe the sequences at the tips of the tree.

Given a tree with *n* species, there are *n −* 1 internal nodes. If each node could be any of the 20 amino acids, there are 20(*n−*1) possible combinations of ancestral states. For just 10 species, one would have to sum over 209 = 512000000000 scenarios; for 40 species, the number of scenarios exceeds the number of atoms in the universe.

Felsenstein realised that not every state needs to be calculated for every global scenario, but an exact calculation can be done through dynamic programming. At each node a *partial likelihood* is calculated representing the (partial) probability of seeing a site conditioned on every observed tip state of tips below that node. By pruning the tree from the tips to the root, a partial root probability can be calculated, which combined with stationary frequencies, provides a site’s probability.

The model assumes each site in a DNA alignment evolves independently. We calculate the likelihood for one site and then multiply the results for all sites.

- At the leaves of the tree, the state is known. If a leaf has an ‘A’, the probability of it being ‘A’ is 1.0, and all the other amino acids are 0.0.
- For an internal node *k* with children *i* and *j*, the probability of node *k* having state *x* is calculated by looking at its children. Just sum the probabilities of all possible transitions from *x* to whatever the children might be.

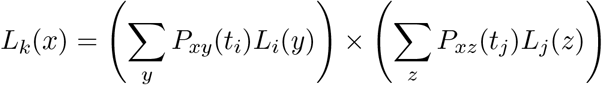

Where *Lk*(*x*) the partial probability of being in state *x* at site *k*, and *Pxy*(*t*) is the probability of state *x* changing to *y* over time *t* (defined by a substitution model like WAG or JTT).
- Once the root is reached, we have the conditional likelihood for each of the 21 states. To get the final likelihood for that site, we multiply these by the prior probability of each state (the stationary distribution) and sum them up. The total likelihood of the observed data at site *i*, given the tree *T* and the substitution model *θ*, is:

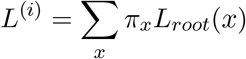

where*πx* the stationary frequency of state *x*, theprobability that a site would be state *x* at the start of the evolutionary process. *Lroot*(*x*) the conditional likelihood that was passed up to the root from its children during the pruning algorithm. It represents the probability of seeing the observed tips of the tree given that the root was state *x*.

Because each site in the alignment *D* is assumed to evolve independently, the total likelihood for the entire sequence alignment of length *L* is the product of the individual site likelihoods:

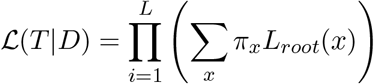

In practice, because multiplying thousands of probabilities results in numbers too small for a computer to handle causing underflow, we use the log-likelihood. This transforms the product into a sum:

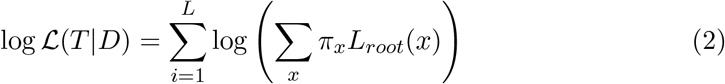

### 2.3 Combining Potts and phylogenetic model

In this paper, we combine the Potts model and Felsenstein’s likelihood by applying the Potts model to the root sequence only, as illustrated in Fig.1. This lets use efficiently calculate the likelihood, while modelling site dependencies in a first order approximation.

**Figure 1.**
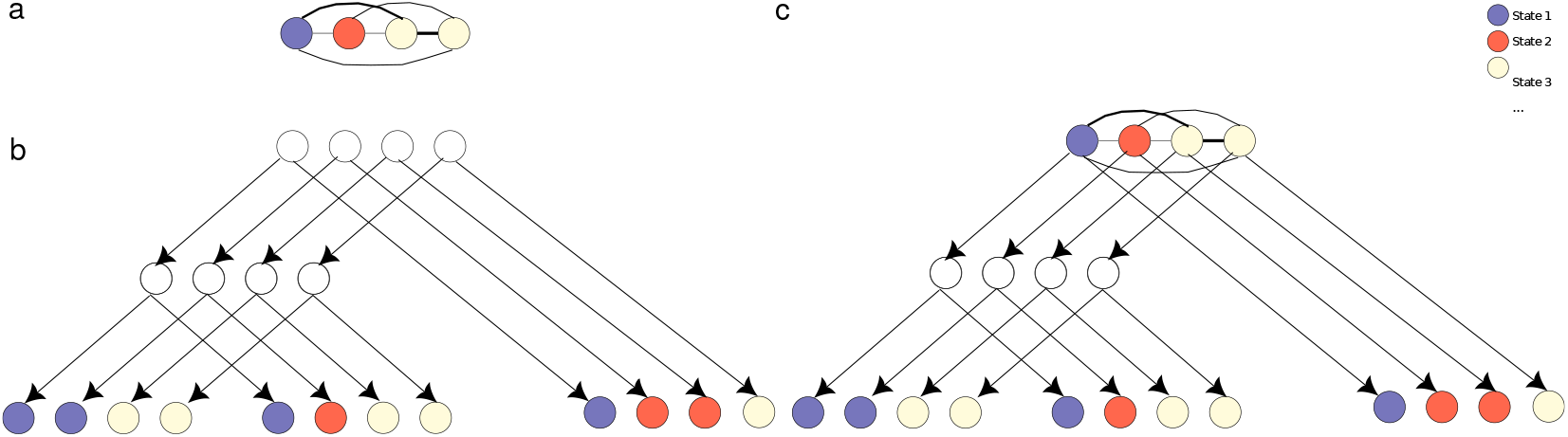
Combining the Potts model and phylogenetic model into a single PhyloPotts model. a) undirected model representing a Potts model: all sites are dependent b) directed graphical model representing a phylogenetic likelihood: all sites are independent, internal node values are integrated out c) combining the Potts and phylogenetic models by applying the Potts model to the root sequence.

Markov chain Monte Carlo (MCMC) is the work horse of Bayesian phylogenetic inference. To use the PhyloPotts model during MCMC, we augment the MCMC state by a sequence *S* representing the root state. This allows us to assing a probability contribution from both the Potts model as well as the tree likelihood. At the root, instead of using the stationary distribution, for each individual site separately, we set the root distribution equal 1 for the state in *S* and 0 for all other states. The joint distribution is the product of the Potts model’s contribution and Felsenstein’s likelihood. Since we are working in log-space, it is the sum of Eq(1) and Eq(2) (but taking root frequencies for each individual site from the root sequence *S*) :

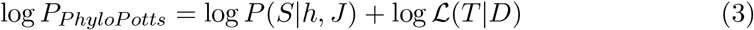

Note that for Potts models, a seperate state representing gaps is used, while in Felsenstein’s likelihood gaps are usually represented as ambiguous characters with a uniform distribution over all non-gap states. Since gaps are important in ancestral state reconstruction, especially when gaps occur in clusters where they are highly dependent on neighbouring states, we use the gap character as a seperate state. This allows us to model the evolution of gaps. Let us define the gap state as the highest numbered state (i.e. state 21 for amino acids). As the 21 by 21 instantaneous rate matrix, we use a standard aminoacid model, like WAG or JTT for the entries *qi,j* where *i <* 21 and *j <* 21 and the rates *q*.*g* and *qg*. represent the rates for mutating in and out of gaps respectively. This results in the following matrix:

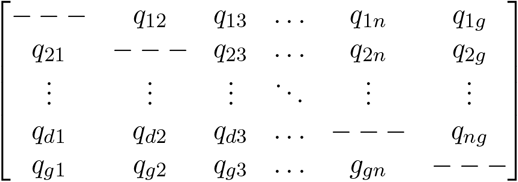

In the experiments that are following, we set the gap rates to constant (*q*.*g* = *qg*. = *c*), but more sophisticated models, like covarion [32] would be interesting to explore.

### 2.4 Simulating from a PhyloPotts model

Seq-Gen[23] is the standard software for simulating the evolution of DNA or protein sequences along a phylogenetic tree. It uses a Monte Carlo approach to evolve sequences from a common ancestor (the root) down to the tips of a user-defined tree. We adapt it to simulate from a PhyloPotts model

The algorithm can be summarised in four main stages:

1. Initialization at the Root: The algorithm begins at the root of the tree by generating a seed sequence of length *L*. In the original algorithm of Seq-Gen, each site in the root sequence is chosen independently by sampling randomly based on the stationary frequencies *pix* defined by the substitution model. To simulate sequence alignments from a PhyloPotts model, we first simulate a sequence from the Potts model by itself, independent of the tree. This can be done by MCMC sampling from Eq(1) alone. A single MCMC proposal is used that randomly picks a site, then randomly proposes a new state for that site. The change in Eq(1) can be recalculated efficiently by only considering the parts that are affected by the new site’s value.
2. Site heterogeneity: Seq-Gen accounts for the fact that some parts of a genome evolve faster than others. A multiplies can be drawn from a gamma distribution, and a site with a high rate multiplier will effectively evolve overe longer branch lengths, increasing its probability of mutation.
3. The Pre-order traversal: The algorithm moves down the tree from the root to the tips. For every branch in the tree, a transition probability matrix *Pxy*(*t*) for that specific branch is calculated using the matrix exponential: 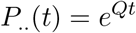

where *Q* is the instantaneous rate matrix defined by the substitution model, containing the relative rates of changing from one state to another, and *t* is the branch length in substitutions per site. For every site in the parent sequence, the algorithm looks at the corresponding row in the *P*..(*t*) matrix to determine the probability of the child site inheriting the same state or mutating to a different one. For parent state *y*, a new state *x* is randomly drawn with probability *P*_*xy*_(*t*).
4. Once the algorithm reaches the tips of the tree, the sequences at those nodes are collected, and these sequences represent the observed data. The internal ancestral sequences are discarded. The final result is a multiple sequence alignment where the patterns of differences between sequences exactly reflect the underlying tree topology and the evolutionary parameters provided.

In contrast to Felsenstein’s likelihood (which works backward from data to find a tree), the simulator works forward from a tree to create data. We will use it to ensure our method is implemented correctly, as it provides a ground truth where the researcher knows the exact history of the sequences.

## 3 Results

To validate the model, we ran a well calibrated simulation study [15] as follows. First we train a Potts model on an existing sequence alignment of 36 aminoacyl-tRNA synthetases (AARSs) for class 2B from [2]. Figure 2 shows a summary of the Potts model inferred from the alignment. The heat map shows that strong interactions are present around the x=y axis, suggesting residues interact highly with neighbours, which is to be expected. However, there are also significant interactions interactions that are non-local, as indicated by the darker patches away from the diagonal, for example, between sites centered around 25, and those centered arond 50. However, there are also some spurious interactions, like between sites 16 and 70. Both of these sites are almost constant (P and R respectively), but show up as highly interacting. We draw 100 ultrametric trees with 50 taxa randomly from a Yule tree prior with birth rate normally distributed (*µ* = 6.0, *σ* = 0.1), giving a range of tree heights from val 0.328 to 1.1198, with 95% highest probability density (HPD) interval 0.3658 to 0.8557 and mean of 0.5976. We use the trained Potts model parameters to simulate sequences alignments based on these trees as described in Section 2.4. Since there are 82 sites in the Potts model, the sequence alignments have 82 sites as well. For the substitution model, we used WAG extended with a gap state as detailed in Section 2.3. Then, we ran BEAST2 on these alignments using the same model and checked whether statistics from the ground truth matched that of the inferred posteriors. For continuous statistics, like tree height and length, the true value should be inside the 95% HPD from 91 to 99 of the 100 runs. For discrete statistics, like the root sequence state, the true amino acid character should be in the 95% smallest coverage set at least 91 of the 100 runs.

**Figure 2.**
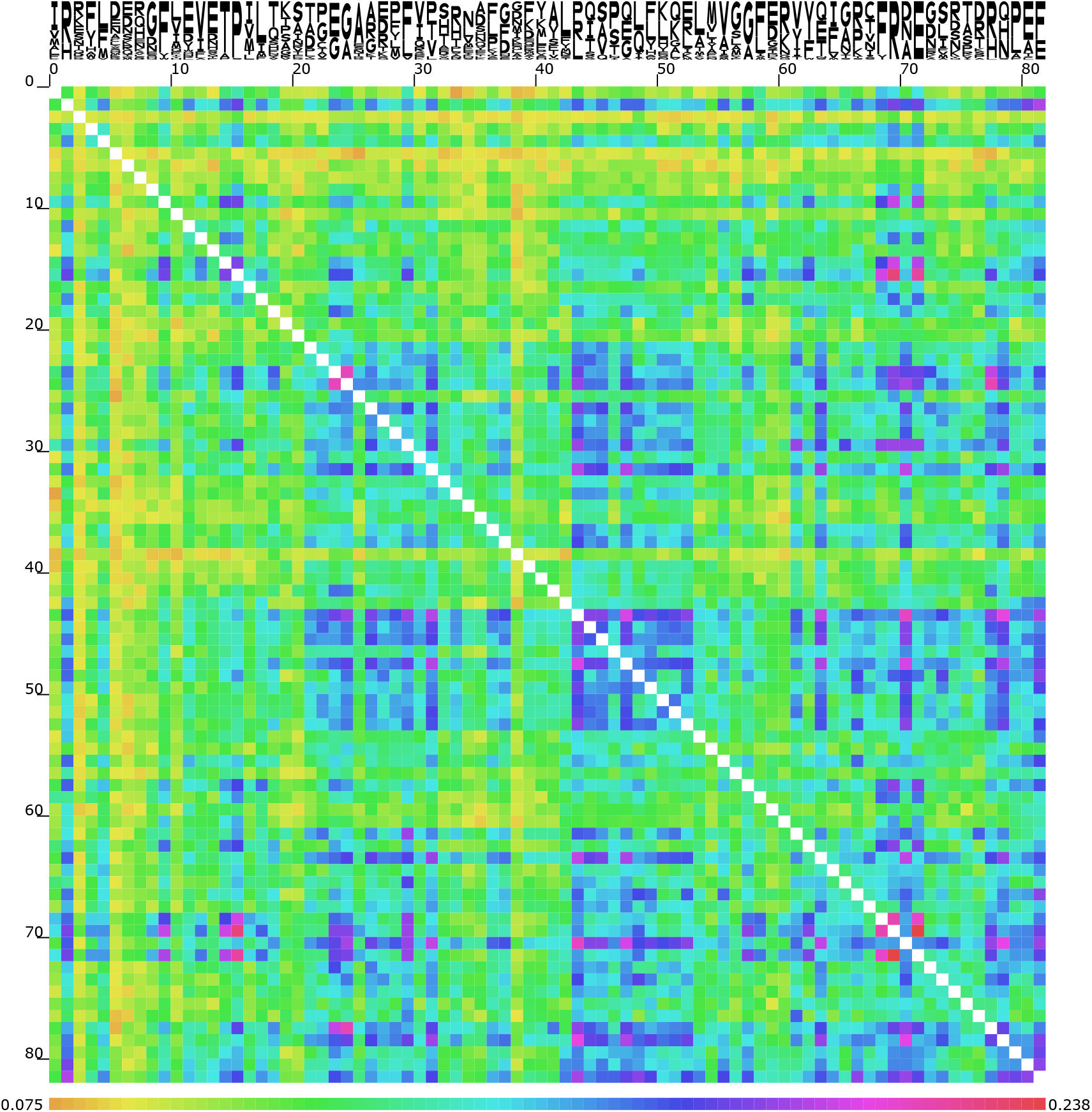
Heat map showing strength of interactions between pairs of sites in the Potts model for the Class 2B sequences. There are 82 sites as indicated by the ticks next to the heat map. The coloured bar at the bottom shows lower interaction at yellow to green and higher interaction at blue to red. Cartoon at the top represent frequencies of amino acids in the alignment by the size of the letters (e.g. site 72 is all E, site 71 just before is about equal parts D, N and A).

Table 1 shows coverage for continuous statistics. It also shows a “triangle” plot [15], where all clade probabilities are collected in bins of size 10% and the bar length shows how often the clade is present in the true tree. Ideally, the bars should all be on the diagonal, but there is some slight variation though no systematic bias. For the root states, coverage is consistently above 95 for all 82 sites. Effective sample sizes (ESS) on average 1822, suggesting good mixing of the root sequence. These results suggest the implementation can infer the original tree correctly.

**Table 1:** Coverage of some statistics in the well calibrated simulation study. The plot shows how often the true tree contains a clade when the inferred posterior is in an interval of size 10% indicated below the plot. The number below at the bottom of the plot is total the number of clades in these intervals. The bars are expected to be close to the diagonal line.

| Statistic | Coverage |
| --- | --- |
| Tree height | 98 |
| Tree length | 97 |
| Yule model | 97 |
| birth rate | 95 |

### 3.1 Sensitivity to Potts model

We repeated the simulation study with the same set of alignments, but doing inference under the standard WAG model with equal frequencies (all amino acids 5% probability, gap states treated as missing data), thus ignoring site dependencies represented by the Potts model. Results for coverage of continuous parameters and clade probability estimates are pretty similar (See Table 2) and even coverage of root state sequence is comparable to that when using the Potts model.

**Table 2:** As Table 1 but inference under independent model.

| Statistic | Coverage |
| --- | --- |
| Tree height | 98 |
| Tree length | 97 |
| Yule model | 97 |
| birth rate | 96 |

However, inspecting the distribution of root states shows significant differences between the models. For example, as shown in Figure 3, though the 95% smallest coverage sets are almost the same, support for different characters is substantially different. This implies ancestral reconstruction between the two models will substantially differ as well.

**Figure 3:**
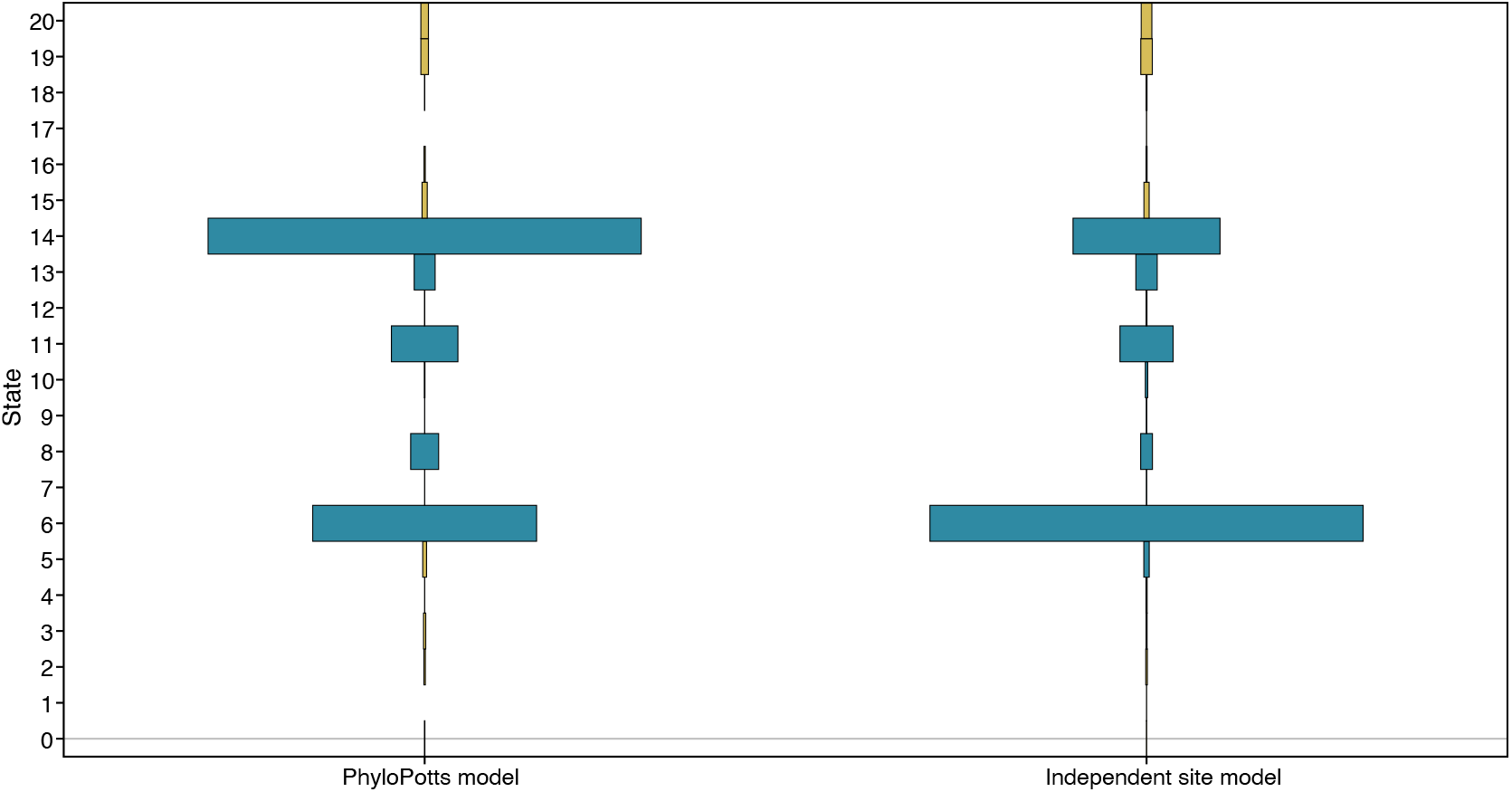
Distribution of a root site sampled under the PhyloPotts model and the standard independent site model. On the y-axis, the 21 states (20 amino acids + 1 gap state) is shown in alphabetical order of their one character representation, and last state is gap state. Width of the bars are proportional to posterior support for the associated amino acid. Blue coloured bars indicate the site is in the 95% smallest overage set, orange it is outside.

The pairs of analyses differ even more when taking correlation between sites in account. Figure 4 shows the correlation between sites 2 and 15 for one of our experimental runs. Appart from the difference in marginal root distributions, the joint distribution is also very different between PhyloPotts and standard model. The PhyloPotts model allows for a lot less uncertainty than the standard model.

**Figure 4:**
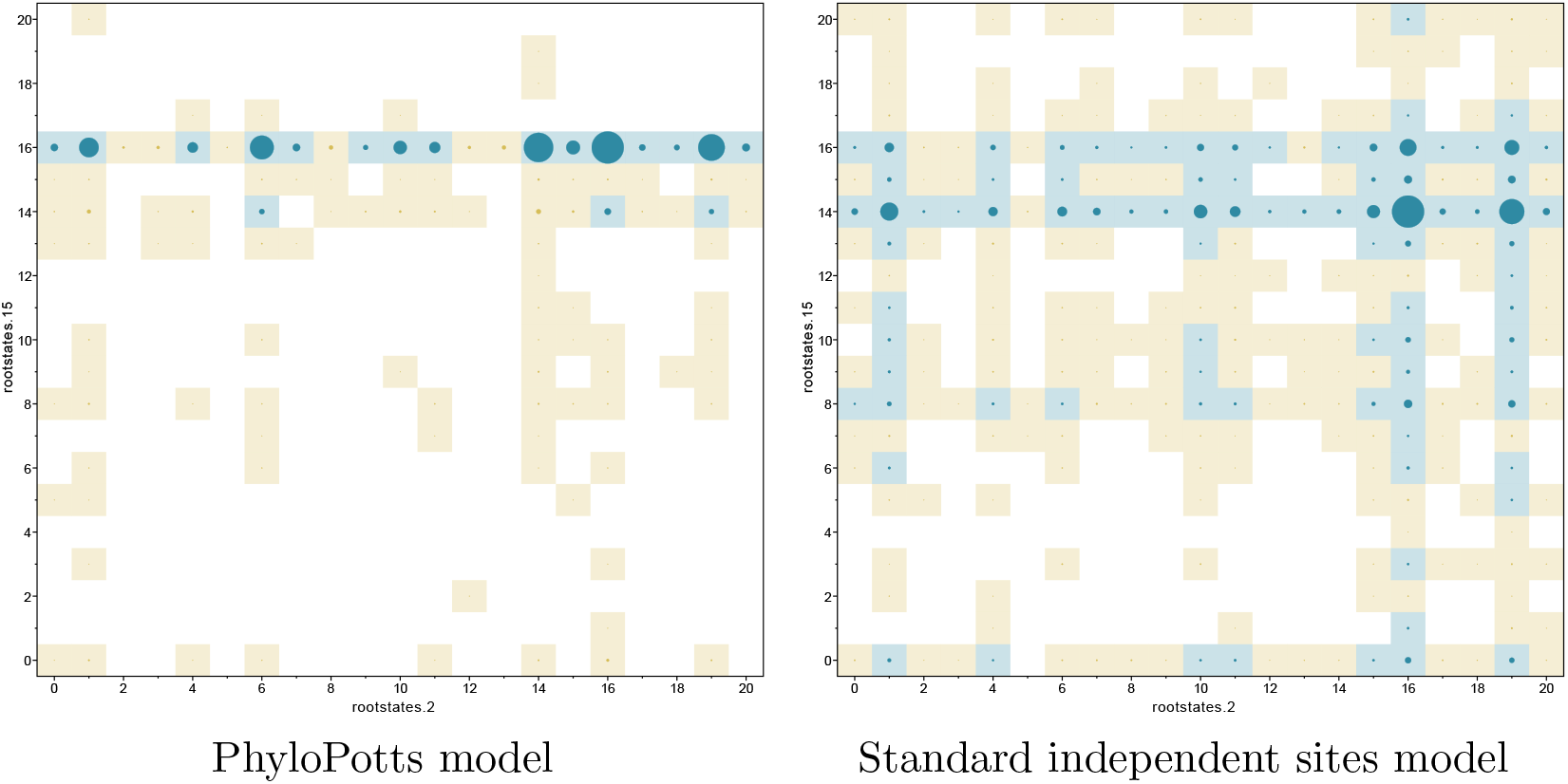
Joint distribution of two root sites sampled under the PhyloPotts model and the standard independent site model. On the x-axis and y-axis, the 21 states (20 amino acids + 1 gap state) are shown in alphabetical order of their one character representation, and last state is gap state. Circle sizes are proportional to joint posterior support for the associated amino acids. Blue colour indicate the combined site is in the 95% smallest coverage set, orange it is outside. There is substantial difference in distribution between PhyloPotts model and independent site model.

## 4 Discussion

The well calibrated simulation study passed with coverage as expected. However, simulating the data, then inferring trees and parameters under the standard independent site model passed coverage of parameters of interest, like tree height and length, as well. Even root state coverage for individual sites is as expected. However, even though 95% smallest coverage sets were satisfactory, considerable differences in root state distributions were observed. Furthermore, comparing pairwise site state distributions also showed considerable differences. From these results, we can conclude that tree topology and node age distributions may not be biased too much by ignoring dependencies between sites. Even though it is known to result in biases[12, 19], for many phylogenetic analyses this may be a minor consideration, thus vindicating standard practice of assuming independence between sites. However, for ancestral state reconstruction we quickly ran into substantial differences when assuming independence (See Figures 3 and 4)

For ancestral reconstruction, consider a part of the sequence alignment that has a high density of gaps. Usually, a sequence will consist of eiter (almost) all gaps, or (almost) no gap at all. Assume that in a 10 site region, 80% of the sequences is all gaps and 20% of the sequences is no gap at all. Standard ancestral sequence reconstruction reconstruct no gaps at all, since gaps are treated as missing data. However, methods that do recognise a seperate gap state, and thus are able to reconstruct gaps tend to have 8 of the sites gap and 2 sites non-gap randomly distributed inside the gap region. This is undesirable, since it is highly unlikely the 2 non-gap sites are plausible to have function in say protein design. Using the PhyloPotts model for ancestral root sequence reconstruction prevents these kind of behaviour: the model will either produce all gaps with 80% probability or all non-gaps with 20% probability.

In this paper, we used a very simple gap mutation model, setting the gap rates to constant. More sophisticated models, in particular covarion models, which allow gaps to be present and/or absent for long periods of time, would be interesting to explore. Another area to explore is the interaction of models for evolution of gaps, like the classical Thorne-Kishino-Felsenstein (TKF91) model[31] with Potts models. Though there are more opportunities for leveraging the power of Potts models for phylogenetics, here only a simple first order approximation of the underlying process was introduced. Capturing dependencies between sites at the level of internal nodes, or even among branches will be computational much more challenging. This will require replacing Felsenstein’s algorithm. A possible solution that looks promising is using explicit mutation annotated trees[13, 17, 20, 33], where instead of integrating out all internal sequence states at the internal nodes in the tree, mutations are sampled on branches of the tree and thus allowing the sequence to be reconstructed anywhere in the tree. This opens up the possibility to combine Potts models at the point where mutations appear, thus allowing more realistic modeling of the underlying process that drives mutations dependent on neighbouring sites.

While for practical reasons, the estimate of Potts model parameters and phylogenetic inference were separated, joint estimation would open a route to then also sample the alignment itself.

## 5 Conclusions

A new model is presented that combines the power of Potts models for capturing structural dependencies with the computational efficiency of Felsenstein’s algorithm for phylogenetic inference. In this paper, we concentrated on protein models, but the PhyloPotts model applies equally well to any alignment of discrete states, like DNA, RNA and 3DI characters. We demonstrated that, though the phylogenetic estimates are not always that less biased compared to using standard models, ancestral reconstruction it significantly impacted by the model.

This paper introduced a proof of concept that combining Potts models and phylogenetics models is possible and desirable. There are a lot of opportunities combining Potts models with phylogenetic inference, including development of richer gap models, allowing more dependencies between sites and joint estimation of Potts model parameters, phylogeny and alignment. Together, this brings has the potential to bring the fields of phylgenetics and structural biology closer together, promising better insights in both fields.

## References

[1] Bouckaert, R., Heled, J., Kühnert, D., Vaughan, T., Wu, C.-H., Xie, D., Suchard, M. A., Rambaut, A., and Drummond, A. (2014). BEAST 2: a software platform for Bayesian evolutionary analysis. PLoS computational biology, 10(4):e1003537.

[2] Carter Jr, C. W., Popinga, A., Bouckaert, R., and Wills, P. R. (2022). Multidimensional phylogenetic metrics identify class i aminoacyl-trna synthetase evolutionary mosaicity and inter-modular coupling. International Journal of Molecular Sciences, 23(3):1520.

[3] Das, R. and Baker, D. (2008). Macromolecular modeling with Rosetta. Annu. Rev. Biochem., 77(1):363–382.

[4] De Leonardis, E., Lutz, B., Ratz, S., Cocco, S., Monasson, R., Schug, A., and Weigt, M. (2015). Direct-coupling analysis of nucleotide coevolution facilitates RNA secondary and tertiary structure prediction. Nucleic acids research, 43(21):10444–10455.

[5] Ekeberg, M., Lövkvist, C., Lan, Y., Weigt, M., and Aurell, E. (2013). Improved contact prediction in proteins: using pseudolikelihoods to infer Potts models. Physical Review E—Statistical, Nonlinear, and Soft Matter Physics, 87(1):012707.

[6] Felsenstein, J. (1985). Phylogenies and the comparative method. The American Naturalist, 125(1):1–15.

[7] Felsenstein, J. (2003). Inferring phylogenies.

[8] Höhna, S., Landis, M. J., Heath, T. A., Boussau, B., Lartillot, N., Moore, B. R., Huelsenbeck, J. P., and Ronquist, F. (2016). RevBayes: Bayesian phylogenetic inference using graphical models and an interactive model-specification language. Systematic biology, 65(4):726–736.

[9] Joy, J. B., Liang, R. H., McCloskey, R. M., Nguyen, T., and Poon, A. F. (2016). Ancestral reconstruction. PLoS computational biology, 12(7):e1004763.

[10] Kamisetty, H., Ovchinnikov, S., and Baker, D. (2013). Assessing the utility of coevolution-based residue–residue contact predictions in a sequence and structure-rich era. Proceedings of the National Academy of Sciences, 110(39):15674–15679.

[11] Khatri, K., Levy, R. M., and Haldane, A. (2025). Phylogenetic corrections and higher-order sequence statistics in protein families: Potts vs multiple sequence alignment transformer machine learning models. Physical Review Research, 7(4):043077.

[12] Larson, G., Thorne, J. L., and Schmidler, S. (2020). Incorporating nearest-neighbor site dependence into protein evolution models. Journal of Computational Biology, 27(3):361–375.

[13] Lartillot, N. (2006). Conjugate Gibbs sampling for Bayesian phylogenetic models. J Comput Biol, 13(10):1701–22.

[14] McGee, F., Hauri, S., Novinger, Q., Vucetic, S., Levy, R. M., Carnevale, V., and Haldane, A. (2021). The generative capacity of probabilistic protein sequence models. Nature communications, 12(1):6302.

[15] Mendes, F. K., Bouckaert, R., Carvalho, L. M., and Drummond, A. J. (2025). How to validate a bayesian evolutionary model. Systematic Biology, 74(1):158– 175.

[16] Minh, B. Q., Schmidt, H. A., Chernomor, O., Schrempf, D., Woodhams, M. D., Von Haeseler, A., and Lanfear, R. (2020). IQ-TREE 2: new models and efficient methods for phylogenetic inference in the genomic era. Molecular biology and evolution, 37(5):1530–1534.

[17] Minin, V. N. and Suchard, M. A. (2008). Fast, accurate and simulation-free stochastic mapping. Philosophical Transactions of the Royal Society B, 363(1512):3985–3995.

[18] Morcos, F., Pagnani, A., Lunt, B., Bertolino, A., Marks, D. S., Sander, C., Zecchina, R., Onuchic, J. N., Hwa, T., and Weigt, M. (2011). Direct-coupling analysis of residue coevolution captures native contacts across many protein families. Proceedings of the National Academy of Sciences, 108(49):E1293–E1301.

[19] Nasrallah, C. A., Mathews, D. H., and Huelsenbeck, J. P. (2011). Quantifying the impact of dependent evolution among sites in phylogenetic inference. Systematic Biology, 60(1):60–73.

[20] Nielsen, R. (2002). Mapping mutations on phylogenies. Systematic Biology, 51(5):729–739.

[21] Pollock, D. D., Taylor, W. R., and Goldman, N. (1999). Coevolving protein residues: maximum likelihood identification and relationship to structure. Journal of molecular biology, 287(1):187–198.

[22] Puente-Lelievre, C., Malik, A. J., Douglas, J., Ascher, D., Baker, M., Allison, J., Poole, A., Lundin, D., Fullmer, M., Bouckert, R., et al. (2023). Tertiary-interaction characters enable fast, model-based structural phylogenetics beyond the twilight zone. bioRxiv, pages 2023–12.

[23] Rambaut, A. and Grass, N. C. (1997). Seq-Gen: an application for the monte carlo simulation of DNA sequence evolution along phylogenetic trees. Bioinformatics, 13(3):235–238.

[24] Robinson, D. M., Jones, D. T., Kishino, H., Goldman, N., and Thorne, J. L. (2003). Protein evolution with dependence among codons due to tertiary structure. Molecular Biology and Evolution, 20(10):1692–1704.

[25] Rodrigue, N., Lartillot, N., Bryant, D., and Philippe, H. (2005). Site inter-dependence attributed to tertiary structure in amino acid sequence evolution. Gene, 347(2):207–217.

[26] Rodriguez Horta, E., Barrat-Charlaix, P., and Weigt, M. (2019). Toward inferring Potts models for phylogenetically correlated sequence data. Entropy, 21(11):1090.

[27] Ronquist, F., Teslenko, M., van der Mark, P., Ayres, D. L., Darling, A., Höhna, S., Larget, B., Liu, L., Suchard, M. A., and Huelsenbeck, J. P. (2012). MrBayes 3.2: efficient Bayesian phylogenetic inference and model choice across a large model space. Syst Biol, 61(3):539–42.

[28] Seidel, S. and Stadler, T. (2022). TiDeTree: a bayesian phylogenetic framework to estimate single-cell trees and population dynamic parameters from genetic lineage tracing data. Proceedings of the Royal Society B, 289(1986):20221844.

[29] Spence, M. A., Kaczmarski, J. A., Saunders, J. W., and Jackson, C. J. (2021). Ancestral sequence reconstruction for protein engineers. Current opinion in structural biology, 69:131–141.

[30] Suchard, M. A., Lemey, P., Baele, G., Ayres, D. L., Drummond, A., and Rambaut, A. (2018). Bayesian phylogenetic and phylodynamic data integration using BEAST 1.10. Virus Evol, 4(1):vey016.

[31] Thorne, J. L., Kishino, H., and Felsenstein, J. (1991). An evolutionary model for maximum likelihood alignment of DNA sequences. Journal of Molecular Evolution, 33(2):114–124.

[32] Tuffley, C. and Steel, M. (1998). Modeling the covarion hypothesis of nucleotide substitution. Mathematical biosciences, 147(1):63–91.

[33] Varilly, P., Schifferli, M., Yang, K., Burcham, T., Cronan, P., Glennon, O., Jacks, O., Laning, E., Marrs, L., Oba, K., et al. (2025). Delphy: scalable, near-real-time bayesian phylogenetics for outbreaks. bioRxiv, pages 2025–03.

[34] Yeang, C.-H. and Haussler, D. (2007). Detecting coevolution in and among protein domains. PLoS computational biology, 3(11):e211.

